# Comparative analysis of miR-155 tissue expression profiles of different breeds of chicken

**DOI:** 10.1101/409623

**Authors:** Sifan Xu, Yang Chang, Guanxian Wu, Wanting Zhang, Chaolai Man

## Abstract

miR-155 is an important microRNA which has multiple functions in many physiological and pathological processes. In this study, partial pri-miR-155 sequences were cloned from AA^+^ broiler, Sanhuang broiler and Hi-Line Brown layer, respectively. Stem-loop RT-qPCR was performed to detect the miR-155 spatiotemporal expression profiles of each chicken breed. The results showed that the partial pri-miR-155 sequences of different breeds of chicken were high conserved. The expression patterns of miR-155 between broiler and layer were basically similar, and miR-155 is expressed highly in immune related tissues. Interestingly, miR-155 expression activity had higher level in fat tissue of the three chicken breeds (14-day-old), but it decreased significantly in fat of the Hi-Line Brown layer (10-month-old and 24-month-old). In addition, the expression activities of miR-155 in 14-day-old broilers (AA^+^ broiler and Sanhuang broiler) were significantly lower than that of Hi-Line Brown layer (14-day-old) (P<0.05). Moreover, miR-155 expression activities in skeletal muscle of 14-day-old and 10-month-old Hi-Line Brown layer were also significantly lower than that of 24-month-old layer (P<0.05). The results indicated that miR-155 might be one of the important factors affecting the differences in skeletal muscle development and adipogenesis between different chicken breeds. These data can serve as a foundation for further study the functions and mechanisms of miR-155 in the physiological and pathological contexts.

## 1. Introduction

MicroRNAs (miRNAs) are a class of endogenous non-coding regulatory RNAs, which are 18-23 nucleotides in length and negatively regulate gene expression at the posttranscriptional levels. miRNAs play critical roles in many biological processes, including cell proliferation, differentiation, development and functions [1-4]. Numerous studies have shown that miR-155 plays important roles in the immune system and can affect the activation of B, T cell and lymphocytes through controlling manifold regulatory roles [5-9]. Interestingly, the emerging evidences of miR-155 role in the development of skeletal muscle and adipogenesis have been noticed recently. On the one hand, miR-155 can repress the expression of MEF2A (a member of the myogenic enhancer factor 2 family of transcription factors) and osteoglycin (Ogn, an important component of the skeletal muscle secretome), and inhibit proliferation and differentiation of myoblast cells [10, 11]. Moreover, miR-155 also regulates the expression of Olfactomedin-like 3 (OLFML3), which may affect prenatal skeletal muscle development in pig [12]. On the other hand, miR-155 can inhibit adipogenesis by directly targeting CCAAT/enhancer-binding protein β (C/EBPβ), cAMP-response element binding protein (CREB) and peroxisome proliferator-activated receptor gamma (PPARγ) [13-15]. In addition, miR-155 also can inhibit brown adipose tissue formation and reduce a brown adipocyte-like phenotype (‘browning’) in white adipocytes in mice. miR-155 and C/EBPβ co-regulate the development of brown and beige fat cells via a bistable circuit [16].

Despite miR-155 has multiple functions in many biological and pathological processes, the possible physiological roles of miR-155 in various tissues between different chicken breeds remain unknown. Now, the poultry industry is facing several problems, including an increase in disease incidence, abundant fat deposits and breed improvement. Therefore, an in-depth analysis of the miR-155 tissue expression profiles in different chicken breeds could provide valuable references to address these issues. It is generally known that there are wide differences on meat quantity and flavor, growth rate and disease resistance between broiler and layer. In this study, three chicken breeds (AA^+^ broiler, Sanhuang broiler and Hi-Line Brown layer) were selected as study objects. AA^+^ chicken and Hi-Line Brown chicken are classical broiler and layer, respectively. Sanhuang chicken, as one breed of broiler, has good meat quantity and flavor, and its production traits are between broiler and layer. We cloned and analyzed partial pri-miR-155 sequences from the three chicken breeds, then analyzed the expression profiles of miR-155 among the three breeds of chicken. We hope the results can provide references for further understanding the functions and mechanisms of miR-155 in chicken.

## 2. Materials and methods

### 2.1 Ethics statement

The proposed study protocol was approved by the Institutional Animal Care and Use Committee (IACUC) of the Harbin Normal University (No.: SYXKHEI2008006). All experiments in chicken were performed in accordance with the Regulations for the Administration of Affairs Concerning Experimental Animals, approved by the State Council of the People’s Republic of China.

### 2.2 Animals and samples collection

Sanhuang broiler (14-day-old), AA^+^ broiler (14-day-old) and different days old Hi-Line Brown layers (14-day-old, 10-month-old and 26-month-old) were all obtained from Harbin local poultry farm. Tissue samples were obtained from the heart, liver, spleen, lung, kidney, thymus, large intestine, small intestine, muscular stomach, glandular stomach, skeletal muscle, skin, brain, fat and bursa of all chicken, then frozen in liquid nitrogen and stored at −80°C.

### 2.3 RNA isolation and cloning of partial pri-miR-155 sequence

Total RNA was isolated from different tissue samples using Trizol reagent (Sigma-Aldrich, St. Louis, MO, USA) according to the manufacturer protocols. RNA samples were digested with DNase I (TaKaRa, Dalian) for 1 h at 37°C to remove genomic DNA. RNA concentrations were measured with ultraviolet spectrophotometer. One microgram total RNA from each sample was reversely transcribed into cDNA using a FSK-100 RT reagent Kit (TOYOBO CO., LTD.) according to the manufacturer instructions.

Partial pri-miR-155 sequences of the three breeds of chicken were cloned using PCR with the cDNA from liver tissue above. The 50 μL reaction system contained 1.0 μL cDNA (20 ng/μL), 4.0μL dNTPs (2.5 mM) (TaKaRa, Dalian), 5 μL 10×Pyrobest buffer II (TaKaRa, Dalian), 2.5 μL 10 pM forward primer (5’-GTGCCCTTAACTTAGACCACATT-3’), 2.5 μL 10 pM reverse primer (5’-TCTAGAGTTCTTCTGTAGGCTGT-3’), 0.5 μL high-fidelity DNA polymerase (TaKaRa, Dalian), and 34.5 μL sterile water. The primers for chicken pri-miR-155 sequence isolation were designed based on the sequence from Gallus (GenBank accession no. DN830517). The PCR program started with a 94°C for 4 min, and 30 cycles of 94 °C /45 s, 56 °C /45 s, 72 °C /30 s, then 72 °C extension for 10 min, finally 4 °C to terminate the reaction. PCR products of the expected size of 263 bp were obtained. The PCR products were cloned into pMD18-T vector (TaKaRa, Dalian) and constructed recombinant vector pMD18-T-155. At least three independent recombinant plasmid clones from each breed were sequenced by Sangon Biotech (Shanghai) Co., Ltd. The partial pri-miR-155 sequences of the three chicken breeds were identical and have been deposited in the GenBank database and assigned GenBank accession no. KU365165.

### 2.4 Homologous analysis

Homologous comparisons of the three partial pri-miR-155 sequences (Hi-Line Brown layer, AA^+^ broiler and Sanhuang broiler) and pre-miR-155 sequences from 23 different species were performed with DNAMAN version 7 software (http://www.lynnon.com), respectively.

### 2.5 Stem-loop real-time quantitative-PCR (RT-qPCR) for tissue expression profile analysis

miR-155 expression in the different tissues was analyzed by stem-loop RT-qPCR. The chicken *U6* gene (GenBank accession no. NR004394) was selected as the internal control. The cDNAs of miR-155 and U6 were produced with the stem-loop primer 5′-CTCAACTGGTGTCGTGGAGTCGGCAATTCAGTTGAGCCCCTATC-3′ and an anchored-oligo (dT)17 primer using the same method as above, respectively.

On aliquots of the cDNA, RT-qPCR was performed simultaneously with miR-155 primers (5′-ACACTCCAGCTGGGTTAATGCTAATCGTGA-3′ (miR-155 forward primer) and 5′-TGGTGTCGTGGAGTCG-3′ (miR-155 reverse primer)) and internal control primers (5′-CTCGCTTCGGCAGCACA-3′ (U6 upstream primer) and 5′-AACGCTTCACGAATTTGCGT-3′ (U6 downstream primer)) with the THUNDERBIRD SYBR® qPCR Mix (TOYOBO), respectively. The dose and reaction procedure of the qPCR reaction system of the miR-155 and U6 were the same. The 20μl reaction system contained 2μl cDNA of each tissue (100ng/μl), 10pM each oligonucleotide primer, 10μl THUNDERBIRD SYBR® qPCR Mix (TOYOBO), 10μl 50X ROX reference dye (TOYOBO) and finally added sterile water to volume 20μl. The PCR program initially started with a 94°C denaturation for 5 min, followed by 40 cycles of 94°C/15 sec and 56°C/45 sec, finally 4°C to terminate the reaction.

### 2.6 Statistical analysis

Three chickens were randomly selected for quantitative analysis in all groups and three technical repetitions per each chicken were carried out. The relative expression activities of miR-155 were calculated using 2^-ΔCt^ method. The data were analyzed by one-way analysis of variance (ANOVA) using IBM SPSS Statistics 20.0 software and GraphPad Prism v5.0 software (GraphPad Software, La Jolla, CA, USA). Differences were considered significant when P<0.05 or P<0.01.

## 3. Results and discussion

### 3.1 Sequence analysis

In this study, partial pri-miR-155 sequences of the three chicken breeds (AA^+^ broiler, Sanhuang broiler and Hi-Line Brown layer) were cloned and sequenced. Homologous comparison of sequencing results revealed that the three partial pri-miR-155 sequences of each breed shared the same length sequence (263bp), and no deletion or insertion event was detected between the different breeds (Fig.1). The flanking ssRNA segments are critical for processing pri-miRNA and the cleavage site is determined mainly by the distance (about 11 bp) from the stem-ssRNA junction [17]. Since the partial pri-miR-155 sequences were identical between the three breeds, we speculated that the mechanisms of processing pri-miR-155 in different chicken breeds were identical.

**Fig. 1.**
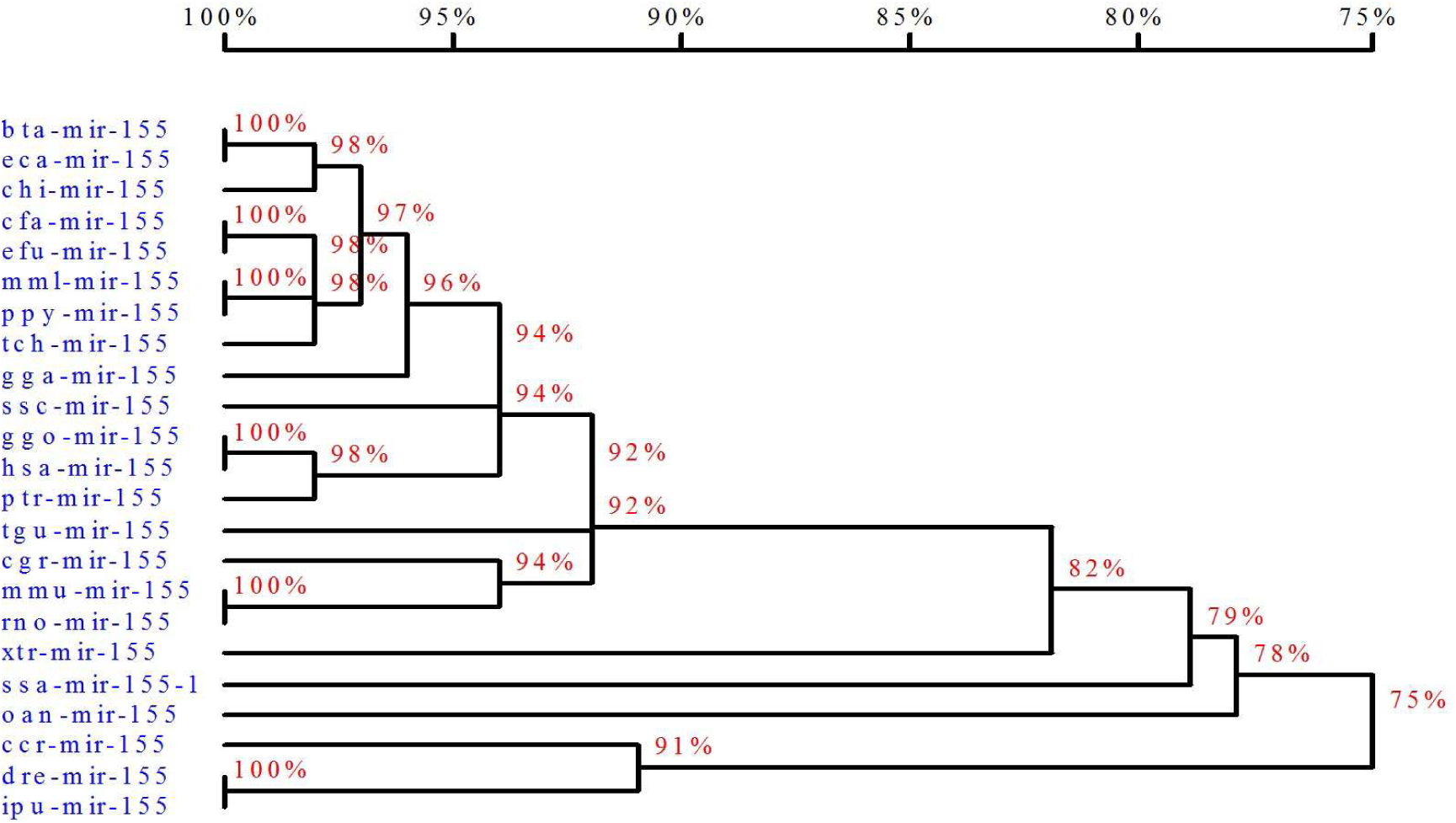
Partial pri-miR-155 sequence of chicken. The sequence of pre-miR-155 is shaded.

As we all know that the sequences of mature miR-155 from different species are highly conservative. However, the homology relationships of pre-miR-155 sequences between 23 different species showed differences (Fig. 2). Homology analysis of pre-miR-155 revealed that chicken pre-miR-155 was most similar to that of land mammals (over 92%) and birds, but not to that of amphibians and fish (below 82%) (Fig. 2). The results implied that chicken should also be a better animal model to study the miR-155 for understanding functions and processing mechanisms among different species.

**Fig. 2.**
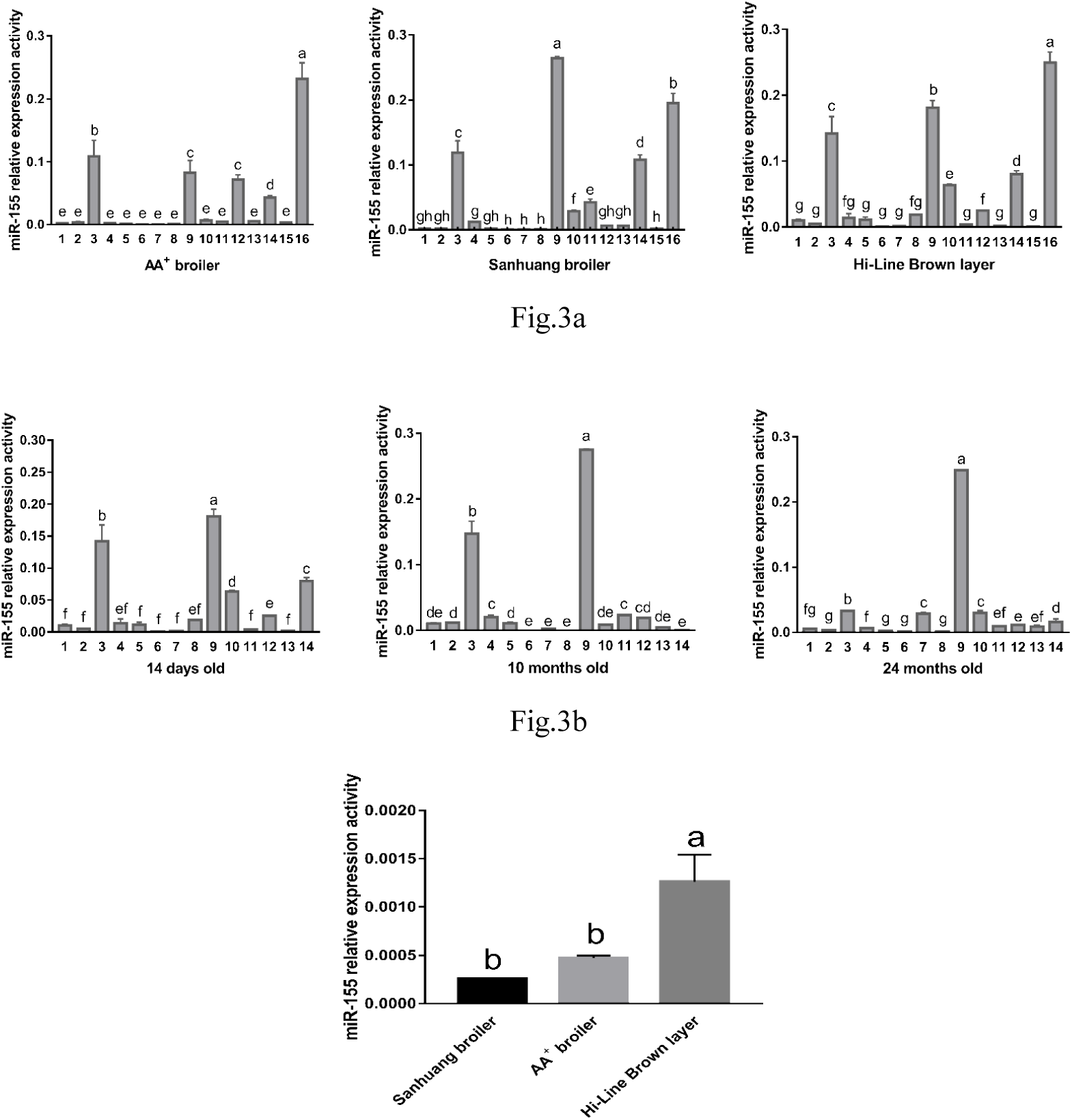
Homology tree of pre-miR-155 based on the nucleotide sequences. aca-mir-155 (Anolis carolinensis, miRBase: MI0018764); bta-mir-155 (Bos taurus, miRBase: MI0009752); ccr-mir-155 (Cyprinus carpio, miRBase: MI0023336); cfa-mir-155 (Canis familiaris, miRBase: MI0008078); cgr-mir-155 (Cricetulus griseus, miRBase: MI0020415); chi-mir-155 (Capra hircus miRBase: MI0030643); dre-mir-155 (Danio rerio, miRBase: MI0002023); eca-mir-155 (Equus caballus, miRBase: MI0012927); efu-mir-155 (Eptesicus fuscus, miRBase: MI0028748); gga-mir-155 (Gallus gallus, miRBase: MI0001176); ggo-mir-155 (Gorilla gorilla, miRBase: MI0020768); hsa-mir-155 (Homo sapiens, miRBase: MI0000681); ipu-mir-155 (Ictalurus punctatus, miRBase: MI0024518); mml-mir-155 (Macaca mulatta, miRBase: MI0007645); mmu-mir-155 (Mus musculus, iRBase: MI0000177); oan-mir-155 (Ornithorhynchus anatinus, miRBase: MI0006775); ppy-mir-155 (Pongo pygmaeus, miRBase: MI0014843); ptr-mir-155 (Pan troglodytes, miRBase: MI0008554); rno-mir-155 (Rattus norvegicus, miRBase: MI0025509); ssa-mir-155-1 (Salmo salar, miRBase: MI0026520); ssc-mir-155 (Sus scrofa, miRBase: MI0015907); tch-mir-155 (Tupaia chinensis, miRBase: MI0031131); tgu-mir-155 (Taeniopygia guttata, miRBase: MI0013842); xtr-mir-155 (Xenopus tropicalis, miRBase: MI0004848); dre-mir-155 (Danio rerio, miRBase: MI0002023). Analysis was done using the DNAMAN software (http://www.lynnon.com). The percentages on the branches represented homology.

### 3.2 RT-qPCR analysis of tissue distribution of miR-155

There are huge differences of growth characteristics between broiler and layer, such as growth speed, egg production and meat flavor, etc. In this study, different chicken breeds were selected to analyze tissue expression profiles, which could provide conveniences for finding the roles of miR-155 in different tissues of chicken. The RT-qPCR results showed that miR-155 was expressed in all tissues tested of the three chicken breeds, but the expression activity differed significantly between different tissues. According to the value of relative expression activity, we divide it into three levels. In AA^+^ broiler (14-day-old), miR-155 was obvious differentially expressed in different tissues, with the strongest expression (the value of relative expression activity is over 0.2) in the bursa; intermediate expression (the value of relative expression activity is between 0.1 and 0.2) in the spleen; low expression (the value of relative expression activity is below 0.1) in the heart, liver, lung, kidney, brain, skeletal muscle, muscular stomach, thymus, skin, small intestine, large intestine, glandular stomach, fat and blood (Fig. 3a). In Sanhuang broiler (14-day-old), the expression profile also presented significant differences, with the strongest expression (the value of relative expression activity is over 0.2) in the thymus; intermediate expression (the value of relative expression activity is between 0.1 and 0.2) in the spleen, bursa and fat; low expression (the value of relative expression activity is below 0.1) in the other tissues (Fig. 3a). In Hi-Line Brown layer (14-day-old), the expression profile also presented significant differences, with the strongest expression (the value of relative expression activity is over 0.2) in the bursa; intermediate expression (the value of relative expression activity is between 0.1 and 0.2) in the spleen and thymus; and low expression (the value of relative expression activity is below 0.1) in the other tissues (Fig. 3a). In Hi-Line Brown layer (10-month-old), the tissues of the strongest and intermediate expression were thymus and spleen, respectively, and other tissues were all the low expression (Fig. 3b). In Hi-Line Brown layer (24-month-old), the tissue of strongest expression was only the thymus, and other tissues were all the low expression (Fig. 3b).

**Fig. 3.**
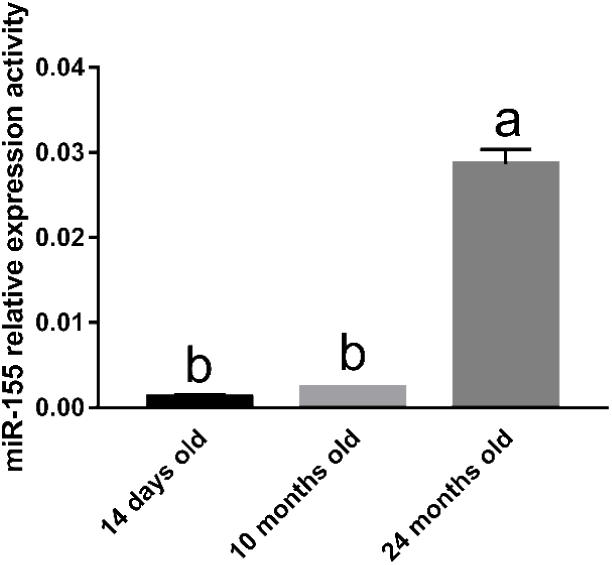
Expression distributions of miR-155 analyzed by stem-loop RT-qPCR. Fig.3a Expression distributions of miR-155 in three breeds of 14-day-old chicken; Fig.3b Expression distributions of miR-155 in different developmental stages of Hi-line Brown layer (14-day-old, 10-month-old and 24-month-old); Fig.3c Expression levels of miR-155 in skeletal muscle of three breeds of 14-day-old chicken; Fig.3d Expression levels of miR-155 in skeletal muscle of different developmental stages of Hi-line Brown layer. 1 heart, 2 liver, 3 spleen, 4 lung, 5 kidney, 6 brain, 7 skeletal muscle, 8 muscular stomach, 9 thymus, 10 skin, 11 small intestine, 12 large intestine, 13 glandular stomach, 14 fat, 15 blood, 16 bursa.

According to the tissue transcription profile analysis of the different breeds of the same age chicken (14-day-old), we found that there was a fundamental similarity of miR-155 relative expression activity in different tissues and organs between the three chicken breeds. For example, miR-155 expression activities were all the strongest expression in the immune-associated tissues (such as bursa, spleen and thymus) between different chicken breeds. As we all know that the spleen, thymus and bursa are the primary lymphoid organs of the chicken immune system. The suitable explanation for this is that miR-155 expression may be related to the biological activities or development of these tissues. However, the differences of miR-155 expression activity in immune-associated tissues between AA^+^ broiler, Sanhuang broiler and Hi-Line Brown layer may be related to the different breeds. Interestingly, we found that the miR-155 expression activity of fat tissue had higher level in the three chicken breeds (Fig. 3a). Studies have shown that miR-155 can inhibit adipogenesis [13-15], whether the miR-155 high expression in fat is related to the less fat deposition of chicken remains to be further studied.

In order to study whether the miR-155 expression is related to development of chicken, the tissue expression profiles were analyzed in different development stages (14-day-old, 10-month-old and 24-month-old) of the same breed (Hi-Line Brown layer). The expression activities of miR-155 in spleen and thymus were worthy of attentions. The expression activities of miR-155 in thymus always kept the strongest expression during different development stages, but the expression activities of miR-155 in spleen decreased significantly in 24-month-old chicken. The possible reason is that the expression of miR-155 maybe related to organ function and characteristics at different developmental stages. It is worth mentioning that the expression activities of miR-155 in fat tissues decreased significantly in 10-month-old and 24-month-old layers (Fig. 3b). Whether it is related to the fat deposition of chicken at these stages also remains to be further studied.

An interesting question that is easily overlooked is that the expression patterns of miR-155 in skeletal muscles of different chicken breeds (14-day-old). Although all miR-155 had low expression activity in skeletal muscle, we found that the expression activities of miR-155 in 14-day-old broilers (AA^+^ broiler and Sanhuang broiler) were significantly lower than that of 14-day-old Hi-Line Brown layer (Fig. 3c). Since miR-155 can inhibit myoblast differentiation and skeletal muscle formation [10, 11], whether the low expression of miR-155 is related to the rapid growth of broilers needs to be further studied in the future.

By comparing the expression activities of miR-155 in skeletal muscle at different developmental stages, it was found that miR-155 had low expression activity in 14-day-old and 10-month-old chickens, but increased in 24-month-old chicken significantly (P<0.05) (Fig. 3d). A reasonable explanation is that the low expression activity of miR-155 can promote skeletal muscle development in 14-day-old and 10-month-old chickens, while the skeletal muscle of 24-month-old chicken has stopped developing, so miR-155 expression activity is increased to inhibit skeletal muscle growth. Regretfully, we do not know whether the strongest expression of miR-155 in the immune tissues is related to chicken immunity status, high expression of miR-155 in the fat is related to chicken fat deposit or meat quantity and flavor, and low expression in muscular tissues is involved in the development of muscle, which are needed to further study in the future.

In conclusion, we cloned the partial pri-miR-155 sequences from broiler and layer, and found the partial pri-miR-155 sequences were high conserved in different breeds. Expression profile analysis of miR-155 showed that layer and broiler had the basically similar expression pattern, and miR-155 was expressed highly in immune related tissues of different breeds of chicken. It worth mentioning is that expression activity of miR-155 shows significant differences in skeletal muscle and fat tissue of different chicken breeds and development stage chicken. The study shows that different-breed chickens are ideal models for study the function and characteristic of miR-155.

## Conflict of interest statement

The authors declare no conflict of interest.

## Acknowledgments

This study was supported by Pre-research project of Harbin Normal University (No. 12XYG-08).

1-gtgcccttaacttagaccacattctaccaagtgcgtctgtttgcagaacaagccatctttttacatgtcacatggaggtcttctcagcgtggcatt gactgattcatgatttctgtcctcctcccacagcagcagtttgttccttgtgagttctgatgagaggcatggtacagtgttaatgctaacatgtagga gtcattcagaggtaaaaacccctatcacgattagcattaacaacatacagcctacagaagaactctaga-263

## Abbreviations

MEF2A: Myogenic enhancer factor 2A
BDNF: Brain-derived neurotrophic factor
OLFML3: Olfactomedin-like 3
Ogn: osteoglycin
C/EBPβ: CCAAT/enhancer-binding protein β
CREB: cAMP-response element binding protein
PPARγ: peroxisome proliferator-activated receptor gamma

## References

[1] Bartel DP. MicroRNAs: Genomics, biogenesis, mechanism, and function. Cell. 2007, 131: 11–29.

[2] Ambros V. The functions of animal microRNAs. Nature. 2004, 431: 350–355.

[3] Krol J, Loedige I, Filipowicz W. The widespread regulation of microRNA biogenesis, function and decay. Nat Rev Genet. 2010, 11: 597–610.

[4] Vigorito E, Kohlhaas S, Lu D, Leyland R. miR-155: an ancient regulator of the immune system. Immunol Rev. 2013, 253(1):146–157.

[5] Xiao C, Rajewsky K. MicroRNA control in the immune system: basic principles. Cell. 2009, 136(1): 26–36.

[6] Vigorito E, Kohlhaas S, Lu D, Leyland R. miR-155: an ancient regulator of the immune system. Immunol Rev. 2013, 253(1):146–157.

[7] Lind EF, Ohashi PS. Mir-155, a central modulator of T-cell responses. Eur J Immunol. 2014, 44(1):11–15.

[8] Jia S, Zhai H, Zhao M. MicroRNAs regulate immune system via multiple targets. Discov Med. 2014, 18(100): 237–247.

[9] Turner M, Vigorito E. Regulation of B- and T-cell differentiation by a single microRNA. Biochem Soc Trans. 2008, 36(Pt 3): 531–533.

[10] Seok HY, Tatsuguchi M, Callis TE, He A, Pu WT, Wang DZ. miR-155 inhibits expression of the MEF2A protein to repress skeletal muscle differentiation. J Biol Chem. 2011, 286(41): 35339 –35346.

[11] Freire PP, Cury SS, de Oliveira G, Fernandez GJ, Moraes LN, da Silva Duran BO, Ferreira JH, Fuziwara CS, Kimura ET, Dal-Pai-Silva M, Carvalho RF. Osteoglycin inhibition by microRNA miR-155 impairs myogenesis. PloS One. 2017, 12(11):e0188464.

[12] Zhao S, Zhang J, Hou X, Zan L, Wang N, Tang Z, Li K. OLFML3 expression is decreased during prenatal muscle development and regulated by microRNA-155 in pigs. Int J Biol Sci. 2012, 8(4): 459–469.

[13] Liu S, Yang Y, Wu J. TNFα-induced up-regulation of miR-155 inhibits adipogenesis by down-regulating earlyadipogenic transcription factors. Biochem Biophys Res Commun. 2011, 414(3): 618–624.

[14] Karkeni E, Astier J, Tourniaire F, El Abed M, Romier B, Gouranton E, Wan L, Borel P, Salles J, Walrand S, Ye J, Landrier JF. Obesity-associated Inflammation Induces microRNA-155 Expression in Adipocytes and Adipose Tissue: Outcome on Adipocyte Function. J Clin Endocrinol Metab. 2016, 101(4):1615–1626.

[15] Rosen ED, Sarraf P, Troy AE, Bradwin G, Moore K, Milstone DS, Spiegelman BM, Mortensen RM. PPAR gamma is required for the differentiation of adipose tissue in vivo and in vitro. Mol Cell. 1999, 4(4):611–617.

[16] Chen Y, Siegel F, Kipschull S, Haas B, Fröhlich H, Meister G, Pfeifer A. miR-155 regulates differentiation of brown and beige adipocytes via a bistable circuit. Nat Commun. 2013, 4:1769.

[17] Han J, Lee Y, Yeom KH, Nam JW, Heo I, Rhee JK, Sohn SY, Cho Y, Zhang BT, Kim VN. Molecular basis for the recognition of primary microRNAs by the Drosha-DGCR8 complex. Cell. 2006, 125(5): 887–901.

